# Effect of cleanup of spiked sludge on corn growth biosorption and leaching of metals

**DOI:** 10.1101/2020.08.21.260661

**Authors:** Driss Barraoui, Jean-François Blais, Michel Labrecque

## Abstract

A chemical leaching process has been used for the cleanup of two municipal biosolids (MOS and BES) spiked with Cd, Cu, Zn or their mixture prior to agricultural use. Non-cleaned, cleaned and washed biosolids were compared as soil amendments for corn cultivation inside glasshouse. Corn growth, biosorption of metals and leaching of these metals in leachate were measured. Results showed that biosolid amendments tend to produce more aerial biomass. Cleanup and washing of BES biosolid significantly augmented total biomass of roots and stalks, respectively. Regarding biosorption of metals, Cd could not be found neither in corn seeds, nor in stalks, while slight amounts of Cu were detected. Whereas Cd and Cu diminished in the order roots > leaves > stalks, Zn diminished from leaves > roots > stalks. Cleanup and washing of MOS and BES biosolids significantly lowered biosorption of Cd, Cu, Zn, and other metals. Leaching into the outlet water varied with time, but average concentrations were moderately low. There were significant amounts of metal leached from MOS biosolid. The effects of cleanup and washing of both biosolids on biosorption and leaching depended on the initial metallic charge and the biosolid type.

## 1 INTRODUCTION

Sewage sludge application improves physico-chemical characteristics of soil, such as organic matter content and water holding capacity, and ensure a similar, or even better yield of plants, as compared to inorganic fertilization [1], [2]. Beneficial effect of sewage sludge can be noticed even two years after their field application, and they also contribute to a stabilization in the grain yield of maize [3]. Indeed, sewage sludge contains important amounts of nutrients that are indispensable to plant growth [4] and it creates very little to zero environmental impact, if utilized properly [5].

One of the main problems related to the agricultural spreading of biosolid is its high content of heavy metals which may be harmful to plants, animals and humans [6]–[8]. Once metals are introduced into lands, they may persist in soil, percolate into leachate or accumulate in plants. Heavy metals can directly or indirectly affect several metabolic processes of plants, like, respiration, photosynthesis, CO_2_-fixation and gas exchange [9], [10]. High application of sewage biosolids could result in heavy metal uptake and many health problems [5].

One of the most problematic metals is Cd. Comparatively, Figlioli et al. [11] reported a higher sensitivity of *Zea mays* to Cd than Pb. It is non-essential and does not have any recognized metabolic role [12]–[14]. Corn is relatively less tolerant to Cd than ryegrass and cabbage, for example [15] (Yang et al., 1996). Yet, An [16] stated that Cd is highly immobilized by roots, and that germination of corn seeds is insensitive to Cd. In contrast, Yang et al. [17] showed that Cd can easily migrate towards corn shoots, and this may explain the ultrastructural damages, especially observed in chloroplasts, induced by Cd [11]. Besides cadmium, Cu is also a widely studied metal. It is an important constituent of many plants’ proteins and enzymes, but a high concentration of Cu may cause chlorosis, inhibition of root growth and damage to plasma membranes [18]. Iron deficiency and chlorosis of leaves are known to be direct consequences of Cu toxicity [19]. Corn is relatively sensitive to Cu excess [18], [20], but is more tolerant than tomatoes [9]. A third important and largely studied metal is Zn. It has a capital role in the activity of many enzymes [21], and is absorbed by plants as the Zn^2+^ ion [22]. Plants use an in-situ complexation of Zn as a strategy to avoid harmful biochemical processes [23]. Interactions such as synergism and antagonism may exist between metals during their uptake in plants [6], [21].

A common way to lower the heavy metal phytoavailability from biosolids is to raise soil pH by adding alkaline amendments for example, by liming biosolids prior to agricultural spreading. However, McBride and Martinez [19] reported that this practice could conjointly causes an increase of the total dissolved organic matter, which in turn could enhance leachability of metals. Biosolid cleanup is then a promising tool to minimize the contamination of soils and waters. Different types of processes (chemical, biological and electrochemical) have been proposed to eliminate toxic metals from municipal sewage sludge [24]–[27].

The present work used a leaching process, called METIX-AC [28], that can diminish the metallic load of municipal biosolids. Two types of biosolids, a non-digested physico-chemical and an aerobically digested biosolids, were sampled from two wastewater treatment plants in the province of Quebec. The biosolid samples were spiked with Cd, Cu, Zn or a mixture of these metals prior to their use as soil enrichment in corn cultivation. The present paper presents results related to the effect of the mentioned biosolid treatments on corn growth on the biosorption of metals in plant parts and on leaching of these metals into the outlet water.

## 2 MATERIAL AND METHODS

### 2.1 Soil and sludge characteristics

The soil used in this study originated from the Montreal Botanical Garden, a loamy-sandy soil (USDA classification) [29] with a high organic matter content. Soil was sieved in order to obtain a uniform grain size (≤ 1 cm diameter). The two municipal sewage sludges tested, were drawn from the province of Quebec, Canada, and included a physico-chemical sewage sludge from the Montreal Urban Community (MOS) wastewater treatment plant (WTP) and a biological sewage sludge from the Haute-Bécancour (BES) WTP. Their nutritional content is presented in Table 1, and amounts of metals supplied by organic amendments are given in Table 2.

**Table 1.** Total solids (T.S.) and concentrations of nutrients (g kg^−1^) in soil and sludge^†^

| Conditions tested | T.S. (%) | TKN | NH<sub>4</sub><sup>+</sup> | NO<sub>3</sub><sup>-</sup> | P | K | Ca | Mg | S |
| --- | --- | --- | --- | --- | --- | --- | --- | --- | --- |
| SOIL | 81.6 | 3.00 | n/a | n/a | 1.04 | 18.0 | 17.8 | 5.72 | 0.32 |
| MOS-CON | 15.6 | 16.3 | 1.40 | 0.04 | 14.6 | 2.63 | 52.7 | 6.41 | 4.77 |
| MOS-CLN | 17.8 | 16.3 | 0.08 ○ | 0.03 | 16.3 | 4.11 | 44.9 ○ | 3.60 ○ | 48.2 ● |
| MOS-WASH | 18.8 | 14.7 | 0.03 | 0.03 | 15.1 | 4.61 | 51.7 | 4.27 | 53.8 |
| MOS-Cd-CON | 14.9 | 14.6 | 1.04 | 0.06 | 12.7 | 6.68 | 61.6 | 7.97 | 6.19 |
| MOS-Cd-CLN | 26.2 | 11.7 ○ | 0.13 ○ | 0.01 ○ | 12.8 | 4.40 | 53.9 ○ | 3.97 ○ | 55.5 ● |
| MOS-Cu-CON | 15.3 | 15.8 | 0.79 | 0.05 | 15.4 | 5.40 | 55.5 | 7.15 | 5.84 |
| MOS-Cu-CLN | 21.8 | 12.0 ○ | 0.10 ○ | 0.03 | 13.7 ○ | 3.98 | 55.7 | 4.22 ○ | 57.1 ● |
| MOS-Zn-CON | 15.3 | 9.80 | 0.74 | 0.04 | 11.5 | 3.84 | 53.8 | 6.64 | 5.16 |
| MOS-Zn-CLN | 21.0 | 11.0 | 0.10 ○ | 0.02 ○ | 10.5 | 4.54 | 53.6 | 4.07 ○ | 58.9 ● |
| MOS-Mix-CON | 16.6 | 8.91 | 0.37 | 0.03 | 9.96 | 3.66 | 58.0 | 7.04 | 6.13 |
| MOS-Mix-CLN | 20.3 | 13.2 ● | 0.11 ○ | 0.03 | 13.8 ● | 2.81 | 48.7 ○ | 3.55 ○ | 52.7 ● |
| MOS-Mix-WASH | 20.4 | 12.0 | 0.03 | 0.02 | 10.5 | 3.46 | 49.7 | 3.78 | 51.8 |
| C.V. (%)‡ | n/d | 6 | 4 | 9 | 5 | 9 | 7 | 7 | 6 |
| BES-CON | 9.78 | 47.2 | 2.96 | 0.10 | 25.0 | 3.24 | 7.69 | 9.54 | 6.46 |
| BES-CLN | 18.5 | 45.8 ○ | 1.11 ○ | 0.11 | 25.0 | 0.72 ○ | 1.05 ○ | 7.04 ○ | 14.5 ● |
| BES-WASH | 18.4 | 51.7 | 0.10 □ | 0.06 | 26.7 | 0.44 | 0.79 □ | 8.00 | 13.9 □ |
| BES-Cd-CON | 8.68 | 30.5 ○ | 2.70 | 0.10 | 18.4 | 3.11 | 7.76 | 9.48 | 6.11 |
| BES-Cd-CLN | 23.7 | 28.6 | 0.23 ○ | 0.03 ○ | 16.9 | 0.70 ○ | 1.17 ○ | 8.74 ○ | 14.1 ● |
| BES-Cu-CON | 8.66 | 32.2 | 3.12 | 0.05 | 16.3 | 3.00 | 8.56 | 11.0 | 6.33 |
| BES-Cu-CLN | 20.6 | 26.6 ○ | 0.26 ○ | 0.03 ○ | 14.4 | 0.69 ○ | 1.19 ○ | 7.91 ○ | 14.3 ● |
| BES-Zn-CON | 9.98 | 32.8 | 4.55 | 0.04 | 17.6 | 2.70 | 7.72 | 10.4 | 5.70 |
| BES-Zn-CLN | 22.9 | 26.7 ○ | 0.26 ○ | 0.04 | 14.0 | 0.54 ○ | 1.10 ○ | 8.18 ○ | 15.1 ● |
| BES-Mix-CON | 8.53 | 31.6 | 2.04 | 0.04 | 13.8 | 2.65 | 7.71 | 10.6 | 5.80 |
| BES-Mix-CLN | 20.4 | 30.3 ○ | 0.35 ○ | 0.03 | 17.9 | 0.72 ○ | 1.13 ○ | 8.19 ○ | 14.7 ● |
| BES-Mix-WASH | 22.9 | 35.0 | 0.13 □ | 0.03 | 22.7 | 0.82 | 0.64 □ | 7.72 | 12.6 □ |
| C.V. (%)‡ | n/d | 6 | 4 | 27 | 11 | 9 | 7 | 5 | 5 |

† Light and dark circles (○,●) indicate respectively significant decreases and increases, due to sludge decontamination. Light squares (□) indicate significant decreases, due to sludge washing. ‡ C.V. (coefficient of variation): to avoid overabundance of data, only an average is given for each element in sludge group.

**Table 2.** Total solids of the maize parts at harvest (%) ^†^

| Conditions tested | Roots | Stalks | Leaves |
| --- | --- | --- | --- |
| SOIL | 50.7 | 28.8 | 81.9 |
| CHEM | 59.4 | 29.3 | 67.8 |
| MOS-CON | 62.3 | 24.5 | 81.4 |
| MOS-CLN | 56.8 | 24.8 | 68.3 |
| MOS-WASH | 62.1 | 19.9 | 85.0 |
| MOS-Cd-CON | 62.2 | 21.1 | 79.2 |
| MOS-Cd-CLN | 53.4 | 23.8 | 80.9 |
| MOS-Cu-CON | 61.5 | 30.3 | 74.1 |
| MOS-Cu-CLN | 58.2 | 20.5 | 82.9 |
| MOS-Zn-CON | 59.0 | 22.0 | 74.0 |
| MOS-Zn-CLN | 62.8 | 27.0 | 77.8 |
| MOS-Mix-CON | 65.4 | 24.5 | 78.3 |
| MOS-Mix-CLN | 57.4 | 26.9 | 74.6 |
| MOS-Mix-WASH | 63.6 | 25.0 | 71.5 |
| BES-CON | 54.7 | 25.2 | 70.9 |
| BES-CLN | 57.8 ● | 18.0 | 82.8 |
| BES-WASH | 60.0 | 29.4 ■ | 76.7 |
| BES-Cd-CON | 57.3 | 23.4 | 75.2 |
| BES-Cd-CLN | 70.1 ● | 25.0 | 84.2 |
| BES-Cu-CON | 60.3 | 24.7 | 82.4 |
| BES-Cu-CLN | 64.6 ● | 22.1 | 78.4 |
| BES-Zn-CON | 64.0 | 18.7 | 80.3 |
| BES-Zn-CLN | 65.3 ● | 25.2 | 82.0 |
| BES-Mix-CON | 48.4 | 15.4 | 85.3 |
| BES-Mix-CLN | 62.0 ● | 20.9 | 76.4 |
| BES-Mix-WASH | 68.2 | 30.7 ■ | 78.5 |

<sup>†</sup> Dark circles (●) and squares (■) indicate significant increasing effects of the decontamination and the decontamination-washing of sludge, respectively.

### 2.2 Sludge spiking procedure

The objective of spiking was to determine whether our leaching process, described below, was able to clean highly loaded biosolids. For this purpose, MOS and BES biosolids were voluntary spiked individually with Cd (as cadmium nitrate), Cu (as copper chloride), Zn (as zinc chloride) or a mixture of these three metals. Additions were performed in such a way as to reach nominal concentrations of about 100, 3 000 and 5 000 mg kg^−1^, respectively of Cd, Cu and Zn in biosolid. Following these additions, and prior to cleanup and washing, biosolids were stored and intermittently stirred during 48 h at ambient temperature, in order to attain redistribution of exogenous metals.

### 2.3 Sludge cleanup and washing

Samples were collected from raw biosolids and were either spiked with metals or left in their original condition. They were subsequently cleaned using the METIC-AC process. Some of these samples were also washed. These non-cleaned biosolid treatments were compared to the cleaned and washed biosolid treatments during the present experiment.

The leaching process consists of adding H_2_SO_4_ and strong oxidants (FeCl_3_ and H_2_O_2_) to leach metals from biosolids. Our leaching process has been experimented in a pilot experiment at the MOS WTP [30]. Extensive details concerning this leaching process are given in the patent [28].

Biosolid washing, which was a supplementary step following the biosolid cleanup, was performed to determine its effect on the elimination of metals that were clustered in the interstitial water of the cleaned biosolid. Due to experimental and laboratory limitations, biosolid washing was not done in the case of individual-metal spiked sludge. It was applied only to the non-spiked, cleaned biosolid and to the fully spiked, cleaned biosolid. Once all biosolids were ready, they were stored in a refrigerated chamber at 4°C for two weeks until they were mixed with soil.

Since the pH of cleaned and cleaned-washed biosolid was too low (pH < 3), so we proceeded to liming of these biosolids before mixing them with soil. This resulted in a neutral pH, which is a condition common to soil and non-cleaned biosolids.

### 2.4 Codification of sludge amendments

The text and the Tables, codes are used to represent the various biosolid amendments: **CON** (non-cleaned), **CLN** (cleaned), **WASH** (washed). For each of the tested biosolids (**MOS** and **BES**), twelve amendments are referred to by:

- **CON, CLN, WASH**: non-cleaned, cleaned and cleaned-washed biosolids, respectively not previously spiked;
- **Cd-CON, Cd-CLN**: non-cleaned and cleaned biosolids spiked with Cd;
- **Cu-CON, Cu-CLN**: non-cleaned and cleaned biosolids spiked with Cu;
- **Zn-CON, Zn-CLN**: non-cleaned and cleaned biosolids spiked with Zn;
- **Mix-CON, Mix-CLN, Mix-WASH**: non-cleaned, cleaned and cleaned-washed biosolids previously spiked with the three metals mixture.

A total of 24 different biosolid amendments of soil were experimented.

### 2.5 Experimental design and soil amendments

The experimental design which was adopted was a randomized block, with a total of 130 pots consisting of 26 treatments replicated five times each. In addition to the 24 biosolid amendments mentioned in the preceding paragraph, two controls were prepared: a non-amended soil (**SOIL**), and a chemically fertilized soil (**CHEM**). The following chemicals were mixed together for the chemical fertilization: NH_4_NO_3_ (34-0-0), Ca(H_2_PO_4_) (0-46-0) and K_2_SO_4_ (0-0-50) to ensure N, P and K requirements, respectively.

In regard to preparation of the culturing medium, plastic pots (29.5 cm diameter, 30 cm height, and a loading capacity of 7-8 kg) were filled with a stratum of gravel enabling leachate to percolate easily. Pots were then completely packed only with soil (in the case of the **“SOIL”** and **“CHEM”** treatments), or only to two thirds of the pot’s height in the case of biosolid amendments. In the latter case, 2 kg of soil-biosolid mixture was spread on the remaining one third height of the pot. An arbitrarily fixed amount of 45 g (dry mass basis) of biosolid was used.

Depending on the nutritional content of each sludge type (Table 1), and in order to avoid differences in the corn growth potentially caused by variable nutrient availability, calculated amounts of inorganic fertilizers, were added to each biosolid amended pot. The same chemicals that were used in the **“CHEM”** treatment were used here. As was recommended by a previous study [31] dealing with the nutrient requirements for an optimum corn growth, exact quantities of the fertilizers were added to the biosolid-soil mixtures, in order to reach the equivalent levels of 160, 110 and 150 kg ha^−1^, respectively of N, P and K.

### 2.6 Cultivation and harvesting of corn

The cultivation of corn was conducted inside controlled glasshouse over a period of fourteen weeks. The G-4011 hybrid of corn was selected, because it is in compliance with the weather conditions of the study zone (thermal needs: 2 500 degrees day). Seeds of corn were sown in plastic pots either in the presence or absence of biosolid. During the first week following sowing, germination of seeds was observed, and twelve weeks later, corn was harvested and divided into its constituents: cobs, stalks, leaves and roots. After weighing each plant part, the materials were oven dried overnight at 105°C to determine dry biomasses. Subsequently, concentrations of P, K, Al, Ca, Cd, Cu, Cr, Fe, Mn, Na, Ni, Pb, S and Zn were quantified in each plant part. The harvested roots were very carefully cleaned from soil, and washed using demineralized water, in an ultrasound bath. This operation was repeated sufficiently to guarantee a dirt-free appearance of roots. Finally, excess water was eliminated with “Kimwipes” paper, and the clean roots were air dried before being weighed and oven dried.

### 2.7 Irrigation scheduling and water sampling

To ensure plant irrigation requirements, demineralized water was delivered by a programmed computer-drip system. Pots containing corn plants were partitioned on three metallic supporting tables. All of the 130 pots were equally and regularly irrigated to maintain soil at the field capacity of water. Hence, sufficient humidity in pots was controlled daily using a manual hydrometer.

A “control drip” was continuously let into a separate container for two purposes: firstly, to check the accuracy of data collected from the previously mentioned computer-drip system regarding the amount of entering water, and secondly to collect a representative sample of irrigation water. Each week, a sample of water was taken from the “control containers” and it was stored at 4°C until analysis.

For the collection of percolating water, a plastic container was hung under each pot of culture. Pots were introduced in polyethylene saucers, which were side-pierced, to allow excess of irrigation water to leach through a pipe inserted into the resulting holes. This handmade system allowed leaching water from each pot to drop into the saucers, then into the hanging plastic containers. Twice a week, leaching water volumes were measured with a graduated cylinder. Following this leaching water quantification, samples of water were stored in small bottles, and conserved immediately at 4^°^C. To constitute a composite sample, aliquots of drained water taken during three succeeding weeks were mixed to the corresponding leaching water volume mentioned above. Given that the total amount of irrigation and leachate was known, it was possible to estimate the part of lost water from each pot during a given period of time.

At the end of each period, the collected composite samples of leaching water and sampled irrigation water were filtered and filtrates were subsequently analyzed for elements mentioned hereafter.

### 2.8 Analytical

Ammonia and nitrate in the biosolid were extracted using a 2 M KCl solution [32] and measured using a Lachat auto-analyzer. After acid digestion, Total Kjeldahl Nitrogen (TKN) was quantified (method: 4500-Nitrogen (organic) B) [33].

The metals content (Al, Cd, Cu, Cr, Fe, Mn, Na, Ni, Pb and Zn) as well as Ca, Mg, P, K and S in biosolid and soil were also acid-digested, before measurement (method No. 30301) [33] by ICP-AES (Varian apparatus, Vista model). Certified samples called RTS-3 (CANMET, Canadian Certified Reference Materials Project (CCRMP)).

Analysis of water pH, was done with a Fisher ACUMET model 915 pH-meter (double-junction Cole-Palmer electrode with a Ag/AgCl reference cell). Ammonium and nitrate were quantified using a LACHAT auto-analyzer (Colorimetric methods QuickChem 10-107-06-2-B (NH_4_^+^) and QuickChem 10-107-04-2-A (NO_3_^−^-NO_2_^−^)).

The concentrations of the following metals Al, Cd, Cu, Cr, Fe, Mn, Na, Ni, Pb and Zn and of the following nutrients Ca, Mg, P, K and S in water and in the different corn parts were determined by atomic absorption with a simultaneous Varian ICP-AES, Vista model. Quality controls were ensured by using certified liquid samples (multi-elements standard, catalog number 900-Q30-002, lot number SC0019251, SCP Science, Lasalle, Quebec) to certify conformity between measurement apparatus.

Detection limits for Cd, Cu and Zn in soil and biosolid were respectively equal to 0.03, 0.12 and 0.18 mg kg^−1^. In the case of leaching water, limits were equal to 0.02, 0.3 and 0.2 μg L^−1^, respectively for Cd, Cu and Zn. The detection limits for Cd, Cu and Zn in the corn tissues were respectively 0.04, 0.27 and 0.09 mg kg^−1^.

### 2.9 Statistical analysis

Analysis of variance (ANOVA) was conducted on all variables. The model measured the importance of the following effects: biosolid type (MOS or BES), spiking with metal(s) (none, +Cd, +Cu, +Zn or +Mix) and treatments (biosolid cleanup and washing). The effect of cleanup was tested for all origins and levels of spiking, but the effect of washing concerned only non-cleaned biosolid samples which were not spiked and those which were fully-spiked (+Mix). Biosolids spiked with an individual metal (Cd, Cu or Zn) were excluded from statistical analysis of the biosolid washing effect, because they were only cleaned, and not washed. Overall, in case that significant effects were obtained, data means were matched using Tukey's HSD. All statistical calculations were executed through SAS (Statistical Analysis Software) version 8 [34].

## 3 RESULTS

Except for the nutritional status of biosolid, the remaining data that are presented and discussed are related specifically to the three heavy metals currently tested (Cd, Cu and Zn) and to some other metals for which concentrations were significantly affected by one or both of the cleanup and washing processes.

### 3.1 Nutritional contents of sludge

The nutrient content of soil and sludge are given in Table 1. The results showed that soil contains high amounts of TKN, K and Ca, but low levels of P, Mg and S. Total solids of tested biosolids are higher in MOS than in BES biosolid, but the latter was richer in all nutritional elements being more concentrated in MOS biosolid. Nitrogen compounds, particularly TKN, are 2-3 times higher in BES biosolid.

The cleanup process has a significant effect on the nutritional content of biosolid. Sulfur augmented in both cleaned biosolids due to the chemicals introduced in the medium during the application of the leaching process. Ca, NH_4_^+^ and Mg were significantly reduced following cleanup of most of the MOS and BES biosolids. Cleanup significantly diminished the levels of TKN, NO_3_^−^ and K in most types of BES biosolid. Meanwhile, TKN diminished only after cleanup of MOS biosolid spiked with Cd or Cu, and augmented with metal mixture-spiking. Finally, the decrease of NO_3_^−^ level in MOS biosolid was related to Cd- and Zn-spiking. In terms of percent of leaching, there were greater losses of NO_3_^−^ and NH_4_^+^ 25-70% and 63-94%, respectively. Similarly, 39-59% of Mg was taken out from cleaned MOS biosolid, while 73-80% of K and 85-86% of Ca was lost from cleaned BES biosolid. In the remaining cases, no more than 26-28% of nutrients leached following the cleanup of one the tested biosolid. Regarding the impact of biosolid washing, there was a significant decrease of NH_4_^+^, Ca and S in the case of BES biosolid.

### 3.2 Production of corn biomass

Data not shown concerning biomass produced a harvest indicated that there was neither a negative effect of biosolid cleanup or biosolid cleanup-washing on the dimensions, nor on weights of corn leaves, cobs, stalks and roots. Moreover, there were generally no significant differences between any biosolid amendment and the soil controls (SOIL). Although biosolid amendments and chemical fertilization (CHEM) generally produced plants with higher aerial parts/roots ratios (fresh weight basis) than SOIL; only MOS biosolid washing had significantly augmented this parameter (data not shown). On the other hand, Table 3 shows that cleanup of MOS biosolid had no significant effect on the total solids of roots, stalks and leaves, whereas significant increases in the total solids of roots and stalks were respectively due to the cleanup and washing of BES biosolid.

**Table 3.** Concentration of tested metals in harvested maize parts (mg kg^−1^) ^†^

| Conditions tested | Cd ‡ |  | Cu |  |  | Zn |  |  |
| --- | --- | --- | --- | --- | --- | --- | --- | --- |
|  | Roots | Leaves | Roots | Stalks | Leaves | Roots | Stalks | Leaves |
| SOIL | 0.08 | 0.06 | 18.2 | 2.16 | 7.06 | 21.3 | 14.0 | 61.0 |
| CHEM | 0.08 | 0.08 | 12.8 | 2.25 | 10.8 | 22.8 | 13.3 | 90.1 |
| MOS-CON | 0.10 | 0.09 | 13.9 | 2.08 | 8.74 | 24.5 | 11.7 | 68.0 |
| MOS-CLN | 0.11 | 0.08 | 12.1 | 2.37 | 7.99 | 22.2 | 13.2 | 58.0 |
| MOS-WASH | 0.10 | 0.08 | 13.2 | 1.48 □ | 9.46 | 27.0 | 12.0 | 78.9 |
| MOS-Cd-CON | 0.54 | 0.26 | 12.4 | 1.48 | 7.03 | 23.2 | 9.41 | 55.3 |
| MOS-Cd-CLN | 0.26 ○ | 0.14 ○ | 12.7 | 1.94 | 10.6 | 32.3 | 9.48 | 62.6 |
| MOS-Cu-CON | 0.07 | 0.06 | 15.8 | 2.65 | 7.80 | 17.9 | 17.1 | 58.5 |
| MOS-Cu-CLN | 0.16 ● | 0.08 | 13.0 | 1.88 | 8.22 | 25.4 | 11.6 | 53.2 |
| MOS-Zn-CON | 0.10 | 0.08 | 10.7 | 1.70 | 7.96 | 36.6 | 10.8 | 54.9 |
| MOS-Zn-CLN | 0.10 | 0.07 | 14.6 | 2.26 | 8.16 | 24.6 ○ | 13.2 | 69.5 |
| MOS-Mix-CON | 0.47 | 0.31 | 14.0 | 1.88 | 7.76 | 30.9 | 12.5 | 73.3 |
| MOS-Mix-CLN | 0.15 ○ | 0.09 ○ | 14.0 | 2.30 | 7.89 | 23.9 | 10.6 | 58.2 |
| MOS-Mix-WASH | 0.15 | 0.08 | 14.1 | 1.67 □ | 10.7 | 24.5 | 10.7 | 57.7 |
| BES-CON | 0.13 | 0.09 | 15.0 | 2.06 | 10.5 | 33.4 | 10.6 | 70.4 |
| BES-CLN | 0.24 | 0.08 | 13.0 ○ | 1.55 | 7.56 | 41.5 | 7.96 | 54.2 |
| BES-WASH | 0.09 | 0.06 | 12.4 | 1.75 | 6.77 | 31.1 | 9.32 | 61.0 |
| BES-Cd-CON | 1.13 | 0.63 | 15.3 | 1.69 | 10.5 | 23.3 | 8.84 | 63.6 |
| BES-Cd-CLN | 0.19 ○ | 0.10 ○ | 13.8 ○ | 2.39 | 10.1 | 29.6 | 16.7 | 75.6 |
| BES-Cu-CON | 0.11 | 0.08 | 15.2 | 2.05 | 9.51 | 24.9 | 10.6 | 59.3 |
| BES-Cu-CLN | 0.13 | 0.08 | 13.4 ○ | 1.83 | 9.92 | 23.8 | 8.40 | 56.4 |
| BES-Zn-CON | 0.16 | 0.08 | 14.2 | 1.39 | 10.5 | 74.9 | 8.20 | 48.8 |
| BES-Zn-CLN | 0.09 ○ | 0.07 | 10.7 ○ | 1.89 | 8.32 | 31.2 ○ | 15.0 ● | 71.8 |
| BES-Mix-CON | 0.95 | 0.47 | 15.8 | 1.31 | 11.2 | 65.7 | 13.7 | 64.5 |
| BES-Mix-CLN | 0.16 ○ | 0.10 ○ | 12.1 ○ | 1.29 | 9.30 | 24.1 ○ | 6.15 | 47.9 |
| BES-Mix-WASH | 0.14 | 0.09 | 9.76 | 3.78 | 20.2 | 24.1 | 13.0 | 73.5 |

<sup>†</sup> Light and dark circles (○,●) indicate respectively significant decreases and increases, due to sludge decontamination. Light squares (□) indicate significant decreases, due to sludge washing. ‡ Cadmium was not detected in stalks and seeds.

### 3.3 Biosorption of tested metals in the corn parts

Data given in Table 4 showed that Cd was generally not detected in stalks, and that in the majority of cases Cd and Cu migrated principally in the roots, leading to the decreasing ranking: roots > leaves > stalks. A different behavior was noted with Zn for which the highest levels in metal concentration were noted in leaves, giving the following decreasing trend: leaves > roots > stalks.

**Table 4.** Concentrations of other metals in harvested roots and leaves (mg kg^−1^) ^†^

| Conditions tested | Roots |  |  |  |  | Leaves |
| --- | --- | --- | --- | --- | --- | --- |
|  | Al | Cr | Fe | Mn | Pb | Pb |
| SOIL | 1990 | 22.1 | 1190 | 60.4 | 0.67 | 2.39 |
| CHEM | 1730 | 30.3 | 1410 | 67.2 | 0.84 | 2.95 |
| MOS-CON | 2070 | 27.9 | 1530 | 61.8 | 1.15 | 2.18 |
| MOS-CLN | 1460 | 16.0 | 1240 | 74.4 ● | 0.77 | 1.65 |
| MOS-WASH | 728 | 11.8 | 791 | 70.4 | 0.70 | 2.67 ■ |
| MOS-Cd-CON | 1270 | 11.8 | 1080 | 57.6 | 0.88 | 3.10 |
| MOS-Cd-CLN | 986 | 18.0 | 1190 | 101 ● | 1.00 | 2.41 |
| MOS-Cu-CON | 942 | 18.5 | 984 | 44.8 | 0.77 | 2.70 |
| MOS-Cu-CLN | 875 | 27.0 | 1070 | 97.0 ● | 0.73 | 2.96 |
| MOS-Zn-CON | 1700 | 19.5 | 1220 | 60.0 | 0.84 | 2.23 |
| MOS-Zn-CLN | 1460 | 23.4 | 1390 | 76.0 ● | 0.82 | 2.47 |
| MOS-Mix-CON | 1140 | 5.91 | 945 | 50.0 | 0.69 | 1.93 |
| MOS-Mix-CLN | 2010 | 24.4 ● | 1490 | 80.0 ● | 1.05 | 2.45 |
| MOS-Mix-WASH | 2320 | 26.8 | 1710 | 79.4 | 1.26 | 1.88 |
| BES-CON | 1160 | 17.1 | 1060 | 99.8 | 0.78 | 1.85 |
| BES-CLN | 847 | 16.0 | 956 | 93.8 | 0.89 | 3.20 |
| BES-WASH | 1680 | 9.15 | 1260 | 94.4 | 1.02 | 2.54 |
| BES-Cd-CON | 719 | 8.62 | 658 | 78.8 | 0.57 | 2.28 |
| BES-Cd-CLN | 2320 ● | 33.7 ● | 1950 ● | 101 | 1.42 ● | 3.29 |
| BES-Cu-CON | 1210 | 29.9 | 1310 | 91.8 | 0.93 | 2.30 |
| BES-Cu-CLN | 1240 ● | 17.1 | 1290 | 105 | 1.05 | 2.37 |
| BES-Zn-CON | 774 | 24.0 | 925 | 105 | 0.72 | 2.76 |
| BES-Zn-CLN | 2270 ● | 36.4 | 1800 ● | 98.8 | 1.08 | 1.93 |
| BES-Mix-CON | 717 | 23.8 | 939 | 80.4 | 0.49 | 2.26 |
| BES-Mix-CLN | 1150 ● | 12.3 | 969 | 74.2 | 0.69 | 1.59 |
| BES-Mix-WASH | 809 | 16.3 | 715 | 75.2 | 0.79 | 2.18 |

<sup>†</sup> Dark circles (●) indicate significant increases, due to sludge decontamination. Dark squares (■) indicate significant increases, due to sludge washing.

Cleanup of MOS and BES biosolids spiked with Cd (alone and in metal mixture) significantly diminished this metal in roots and leaves. Similarly, cleanup significantly diminished Cd in roots in BES biosolid spiked with Zn, while an increase occurred in MOS biosolid spiked with Cu. The Cu and Zn contents in leaves were not affected by the cleanup of any biosolid. With BES biosolid, cleanup significantly diminished Cu in roots. The Zn decrease in roots was recorded following cleanup of biosolid spiked with Zn alone in both MOS and BES biosolids and in with metals mixture in the BES biosolid only. In contrast, Zn significantly augmented in stalks following cleanup of BES biosolid spiked with Zn alone. There was only one case where biosolid washing had a significant effect, namely on Cu in leaves corresponding to MOS biosolid amendments.

### 3.4 Biosorption of other metals in the corn parts

All significant effects of biosolid cleanup noted mostly in BES biosolid were manifested as increases in the concentration of Al, Cr, Fe, Mn and Pb in roots. Washing of MOS biosolid resulted in an increase of the Pb level in leaves (Table 4). MOS biosolid cleanup increased Mn and Cr levels in roots, but the Cr increase was restricted to biosolid previously spiked with the metal mixture. In the case of BES biosolid, cleanup significantly augmented metals in roots, especially Al in all BES biosolid types, Cr and Pb in biosolid spiked with Cd alone and Fe in biosolid spiked individually with Cd and Zn. Finally, non-spiked MOS biosolid washing significantly augmented Pb in leaves.

### 3.5 Lixiviation of metals through leachate

The average concentrations of metals in leachate (Table 5) were calculated from data corresponding to the four consecutive periods (P1 to P4). All volumes of the leachate samples have been taken into consideration for the calculation of the average concentrations of metals in solution. Statistical analyses of data were performed on raw data for each separate period. When a significant effect of biosolid cleanup was noted during a given period, it was reported in Table 5 by specifying, between brackets, the period of occurrence.

**Table 5.**
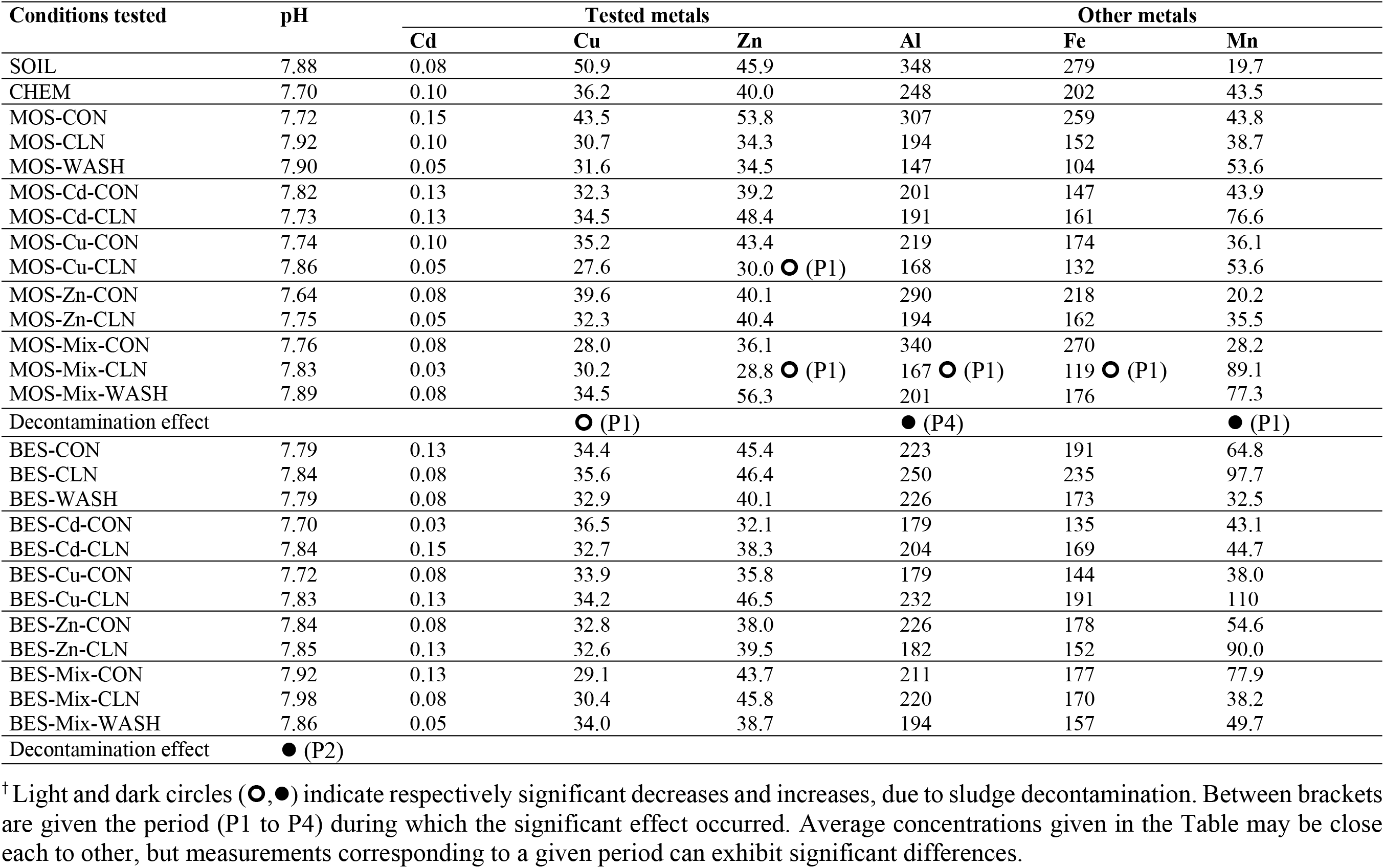
Average concentrations of tested and other metals leached in drainage water (μg L-1) ^†^

It is shown in Table 5 that the average concentrations of metals which leach from biosolid amended pots were lower (Al, Cu and Fe) or higher (Mn and Zn) than those of non amended pots, while approximately similar results were noted in the case of Cd. The pH of leachate samples averaged between 7.64 and 7.98.

Table 5 indicates that no significant effects of washing were observed for both biosolids, whereas cleanup affected the metals leaching only from MOS biosolid amended pots. In the case of BES biosolid, only leachate pH significantly augmented from 7.66 to 7.99 during the second period, P2 (data not shown). In the case of MOS biosolid, changes were mostly noted during the first period, P1, where MOS biosolid cleanup led to a decrease of Cu in leachate for all biosolid types, of Zn for biosolid spiked with Cu, of Zn and metal mixture as well as Al and Fe for sludge spiked with the metal mixture. In contrast, MOS biosolid cleanup significantly augmented the concentration of Al in leachate during the last period, P4, and of Mn during the first period, P1.

Leaching of metals over the time was also studied (data not presented). It was shown that for all treatments, leaching of Cu and Zn were similar in that they were high at the beginning of the experiment, and then gradually diminished with time. An inconsistent decrease in concentration was noted for Cd at the beginning of the experiment, and for Cu and Zn at the end of the experiment.

## 4 DISCUSSION

The leaching process diminished concentrations of heavy metals in biosolids, even when high loads of metals are present. The present data show that although cleanup diminished the nutritional content of MOS and BES biosolids, there were still satisfactory levels of many nutrients allowing the profitable reuse of these biosolids as soil amendments. Cleanup of both raw biosolids (non-spiked) generates biosolids containing higher levels of nutrients than those desired for an agricultural spreading of biosolid. These levels (g kg^−1^, dry mass basis) were satisfactorily reached following cleanup of BES biosolid (TKN > 27, NO_3_^−^> 0.01, P > 18 and Mg > 5.6) as well as that of MOS biosolid (NH_4_^+^> 2, K > 2.5 and Ca > 19). Washing of both biosolids had few or even no significant effects on the concentration of nutrients, compared to the impact of cleanup.

Among the subsequent effects of the application of sludge containing high levels of metals are the risk of provoking a limited growth of plants [6], an excessive biosorption of metals which may be transmitted throughout the food chain [14] or even an occurrence of toxicity [35].

Regarding the plants’ development, although not shown, our results indicated that nor the dimensions, nor weights of corn leaves, cobs, stalks and roots were negatively affected by sludge treatment. These data are in concordance with those of Szymańska et al. [3], where biosolid fertilization, versus mineral fertilization, of corn grown for grain, did not reveal any difference regarding the growth and the development of plants.

It can also be stated that in our case, even when spiking MOS and BES biosolids with Cd, Cu and/or Zn, no perceptible results were noted in the germination rate of seeds, nor in any plant growth parameter, with only few exceptions. The effects of cleanup and washing on biosolid were observed mainly as an increase in the aerial parts/roots ratio of weights in the case of cleanup of MOS biosolid or in the total solids of roots and stalks, respectively following cleanup and washing of BES biosolid. The absence of important differences between any biosolid amendment and SOIL concerning the growth performance of corn plants might be explained by the small amount of sludge that was tested (45 g pot^−1^: the equivalent of about 7 t ha^−1^). Even if the sludge rates used in our experiments are much higher, Szymańska et al. [3] obtained similar results, although they considered an application of 10 t dry mater per ha^−1^ of biosolid once every five years (mean of 2 t ha^−1^), during a corn biosolid amended study in the field conditions. To obtain significant differences compared to unfertilized soil, one has to consider higher sludge application. Ilie et al. [4] observed during an experimental pots study, that sewage sludge fertilization produces significant increase of corn yield that is evident starting with 200 kg N ha^−1^ rate (equivalent to 10 t ha^−1^), compared to the soil control where the lowest yield was obtained. Furthermore, Tejada et al. [36] obtained 17% increase in the corn yield, with the foliar application of sewage sludge, compared to the untreated samples.

The lack of significant differences we noticed was probably influenced by the rich character of the soil. Non-cleaned and cleaned biosolids have strongly similar results in terms of detailed corn growth data. As mentioned earlier, it appears that biosolid is nutritionally rich so that treatment leaves it still capable of ensuring adequate development of corn plants. Greater differences between controls and biosolid amendments as well as between cleaned and non-cleaned biosolids could probably be expected if one or both of the following conditions were encountered: the soil tested did not contain such high concentrations of nutrients, and the biosolid samples were larger than 45 g pot^−1^.

With respect to the uptake and biosorption of metals, it was shown that in all corn parts, Zn was more concentrated than Cu which, in turn, was more abundant than Cd. The latter was undetected in seeds and in stalks, but it was more highly concentrated in roots than in leaves giving a shoot/root ratio equal to 0.33. Cu showed a similar trend as that of Cd, but with a clearer distinction between the metal levels in roots versus in leaves. The lowest Cu amount was retrieved in stalks. Data showed that the metal that mostly stored in plants was Zn. Leaves had the highest concentration of this metal, followed by roots and stalks. These results indicated that Zn seemed to be the most mobile metal since it appeared to be more easily transferred to aerial parts. Cd and Cu showed an opposing trend, as they were mainly confined to roots. The remarkably higher uptake of Cu by roots, along with a limited translocation to shoots, agrees with previous works (Ouzounidou et al. 1995; Mocquot et al. 1996; McBride 2001; Ali et al. 2002).

The mobility of Zn toward aerial parts of corn was confirmed by Ilie et al. [37], who showed that sewage sludge rates higher than 300 kg N ha^−1^ (equivalent to 15 t ha^−1^ of sludge) resulted in statistically increases of Zn content in the corn kernels.

When biosolid was spiked with a given metal its concentration augmented in one or more parts of corn plants, but this increase was very limited in the case of Cu. The use of Cd-spiked MOS biosolid enhanced Cd absorption in roots and leaves, respectively by 5 and 3 times. Similar results were obtained when BES biosolid was spiked with Cd and with the metal mixture. In the case of Cd spiking, the uptake of Cd was augmented by 9 and 7 times, while the metal mixture caused an increase in Cd uptake by 7 and 5 times, in roots and leaves, respectively. Such results, showing that Cd uptake is augmented by its level in the medium are supported by the work of Ilie et al. [37], who reported that Cd level in the corn leaves increased directly proportional with the rate applied of sewage sludge. Also, following the use of Zn-spiked BES biosolid, metal uptake by roots was doubled. Thus, it is well established that as the total concentration of a metal in the soil increases, so does the probability of its uptake by roots. However, metal speciation is of great importance for judging the uptake of metals by roots as well as their possible biosorption in plant organs [38].

The effects of biosolid cleanup on the uptake and biosorption of the tested metals were far more significant on BES biosolid than on MOS biosolid. Cleanup can counter the spiking-induced increase in absorption of metals. To be more specific, uptake of Cd and Zn by roots can be reduced by up to 84% and migration towards leaves can be reduced to 63%. These results pertain to BES biosolid. Although spiking biosolid with Cu (alone or in the metal mixture) did not provoke an increase of Cu concentration in a given corn part, cleanup of BES biosolid caused a decrease in the metal’s uptake by roots by about 10-25%. Overall, it can be stated that our leaching process significantly diminished the levels of Cd, Cu and Zn in both biosolids to such levels that their uptake by roots, and consequently their transfer in a given corn tissue, can be greatly diminished. Moreover, cleanup of BES biosolid spiked with Zn led to a decrease in the uptake of Cd and Zn by the roots. Controversy exists, however, in the published results regarding the competition that might exist between metals for their uptake by plants, suggesting that Zn may compete with Cd, since they use the same transport site [6] or that alleviation of Cd toxicity by application of Zn is difficult to assess, because of the implication of nutrients such as P [14].

More recently, the work of Przygocka-Cyna and Grzebisz [39] showed that any increase in Fe concentration in corn grain resulted in a simultaneous decrease in Cd concentration, attesting then for an antagonism behaviour between these two elements. More clearly, Przygocka-Cyna and Grzebisz [39] stated that an increased exogenous supply of Fe results in decreased uptake by plant roots. And this indeed the case with our experiments, where the concentration of Fe was significantly augmented in both cleaned biosolids by 13% to 20% as this metal is added (as ferric chloride) to biosolid during the application of the leaching process [28].

Therefore, the augmentation observed in the availability of Cd and Zn due to spiking reflects an increase of their uptake by roots and their translocation to leaves (Table 3). However, cleanup-washing of BES biosolid diminished the proportion of the available fractions, which reduced their content in the corn tissue. The increase noted for the available fractions of metals in the cleaned-washed MOS biosolid did not reflect an additional biosorption in corn plants. The reason is probably that there was competition for the uptake with other metals such as Mn, which augmented following the cleanup-washing of all MOS biosolid types or Fe which augmented following the cleanup-washing of several BES biosolid (Table 4) reflecting the antagonism process which could occur between Cd and Fe, as this was already explained above.

An analysis was done on the toxicity of cadmium, copper and zinc, and included a comparison of experimental data obtained in this project with published results [40]. Indeed, one of the previous experiments, dealing with the effect of our leaching process on the toxicity and bioavailability of metals showed that the biosolid treatment significantly diminished the biosorption of cadmium, copper, and zinc in several exposed plant species [40]. These findings are then similar to those of Al-Busaidi and Mushtaque [5], who showed that no toxicity, neither excessive biosorption of heavy metals occurred when sewage sludge are treated with green wastes (Kala compost) to serve as soil amendment for plants’ cultivation.

Empirical data did not show any significant decreases in root length and biomass production that betray Cu toxicity [16], [18], [41], [42]. Dry biomass yield of corn was not affected by Zn [43]. Similarly, Cd seemed not to cause a decrease in root and shoot biomasses [15], [43] and no nutritional deficiency was noted [13]. Lagriffoul et al. [44] however, stated that variation in growth and in mineral content of Cd-contaminated corn seedlings are not direct consequences of Cd uptake by plants. Oppositely, Figlioli et al. [11] stated that Cd induces ultrastructural damages, which are especially observed in chloroplasts. Contradictory statements reported in the literature concerning the effect of heavy metals on the development of plants are certainly due to differences in the experimental conditions. Currently, the absence of toxic effects can also be linked to the small quantity of biosolid tested as well as to the fact that bioavailable forms of metals, such as exchangeable fractions, were extremely low, both before and after biosolid cleanup and washing. In this context, it is well known that availability of biosolid-borne metals in soils is relatively low because these elements are immobilized mainly as oxides [21].

The existence or absence of metal toxicity in the current experiment can also be explained by comparing recorded data to the published toxicity thresholds. First, Cd level in any corn part was far below the toxicity limits of 4-30 mg kg^−1^ [45], [46]. The phytotoxicity of Cu is very well documented but there are disagreements concerning the toxic threshold of this metal in plant tissues. A large range (20 to 100 mg kg^−1^) was generally fixed for plants [45], [46], but in the specific case of corn, Fageria [47] stated that if the Cu level in soil reaches 48 mg kg^−1^, metal concentration in plant tissues attains the toxic onset of 11 mg kg^−1^. Yet, Mocquot et al. [9] and Borkert et al. [20] reported that Cu toxicity to corn occurs only when metal concentration in leaf or root tissues reaches 20-21 mg kg^−1^. McBride [38] criticized the USEPA risk assessment which assumes that in biosolid amended soils, Cu toxicity to corn does not occur even when metal concentration in shoots reaches 40 mg kg^−1^. In our case, Cu toxicity seemed not to occur in the presence of sludge. The highest Cu concentration was noted in roots (18.2 mg kg^−1^) for SOIL, which initially contained 23.4 mg Cu kg^−1^. However, all biosolid amendments diminished the Cu concentration in roots, as compared to SOIL, by up to 41%. In the case of Zn, whatever the treatment of concern, the metal concentration in all corn parts was far below the toxicity limits of 400-1000 mg kg^−1^ [45], [46].

Regarding the remaining heavy metals measured in corn parts (data not shown) indicated that Mn migrated mostly towards leaves, but it was below 400 mg kg^−1^, far below the toxic level of 2 480 mg kg^−1^ [47]. Comparatively, Ilie et al. [37] showed that Mn augmented in leaves, in parallel with the increase of biosolid quantities in pots, while it diminished in the corn Kernels. Similar to the results reported by Ali et al. [18], Pb had stored more in leaves than in roots, but metal concentration was below the toxicity limits of 30-300 mg kg^−1^, whereas Cr sometimes exceeded 4-8 mg kg^−1^ [21], [46]. To explain the behavior of Pb, we could mention the work of Figlioli et al. [11], who reported the complex pattern of Pb uptake by the corn parts when Pb enters the plant roots from Pb-enriched soils, it shows small translocation to the aerial parts; nonetheless, increased concentrations of Pb in aboveground tissues can be caused by entering of metal-bound dust and fine soil particles, or directly to leaves through stomata [11].

In leachate, no clear effect of BES biosolid cleanup was noted, except for a significant increase in pH. The concentrations of Al, Cd, Cu, Fe, Mn and Zn seemed unaffected when MOS and BES biosolids were washed. The significant increase in pH may have been caused by a large increase of NH_4_^+^ ions in cleaned BES biosolid (data not shown).

The highest concentrations of Cd, Cu and Zn in leachates that issued from biosolid amended pots were respectively 0.15, 43.5 and 56.3 μg L^−1^. In comparison, Gray et al. [48] measured the concentration of Cd in leachate from New Zealand pasture soils that were treated repetitively with phosphate fertilizer, and found higher results: 0.35-1.57 μg Cd L^−1^. Li et al. [49] tested mixtures of soil with unstabilized or stabilized sludge (using phosphorus based products), and measured metals (Cu, Cr and Zn) leaching in the outlet water. Total inputs of metals were lower (in case of Cu and Cr) or relatively close (in case of Zn) to our tested loads. Although, the levels of all three metals, Cu, Cr and Zn, in leachates reported by Li et al. [49] attained concentrations as high as (in μg L^−1^) 1 340, 70 and 1 060, respectively. However, one has to take into account that both experiments are quite different each from other. Li et al. [49] did not consider any plant uptake, but only leaching of metals from cylinders filled with biosolid-soil mixtures, that were irrigated with distilled water. Nevertheless, it should be highlighted the importance of our leaching process, which ensures removal of the most labile fractions of metals from biosolids, prior to soil amendments. Cleanup of biosolids as well as corn plant cultivation may explain the large gap observed between our results and those of other researchers.

Although this is far beyond the scope of the present work, the metal content of the sampled leachates can be compared to a severe criterion such as the guidelines for drinking water quality, in order to evaluate the risk of groundwater contamination by these leachates. Data showed that average concentrations of most metals in leachate were far below the drinking water quality guidelines of the Canadian government, of the World Health Organization (WHO) and of the European Economic Community (EEC). This was particularly the case for tested metals (Cd, Cu and Zn), for which averaged concentrations in leachates did not exceed 5, 1 000 and 5 000 μg L^−1^, respectively. The Fe concentration was also far below its safety guideline of 300 μg L^−1^ [50]. The mean concentrations of Fe and Zn were likewise below the respective toxic thresholds of 300 and 180 μg L^−1^, concentrations at which these metals are detrimental to the freshwater fish [51]. Such was not the case of Al and Mn which exceeded their respective limits of 200 and 50 μg L^−1^ [50]. Al content in drinking water is of great concern, because a close correlation seems to link the amount of metal that is consumed by humans and the occurrence of Alzheimer’s disease [52]. In this project, the highest Al concentration (348 μg L^−1^) was obtained for leachates of non-amended pots, but all biosolid amendments diminished this metal level by up to 58%. On the other hand, data not shown indicated that Pb was not detected in leachate, and Cr did not exceed the critical value of 50 μg L^−1^ [50]. Finally, data not shown indicated that leaching of the three tested metals (Cd, Cu and Zn) tend to decrease, with time leading to lower concentrations at the end of experiment. Time variation of leaching fluctuated more in the case of Cd than Cu or Zn. It can thus be stated that the quality of water that leached from pots amended with cleaned biosolid is certainly of acceptable quality to be used for agricultural irrigation purposes.

The differences observed between MOS and BES biosolids in terms of biosorption and leaching of metals are probably caused by metal speciation in tested biosolids which were differently affected by our leaching process. However, the speciation and consequent bioavailability of metals depend on many soil parameters, among which the pH is of great importance. For example, application of organic matter to an acidic topsoil increases the Zn part that is held conjointly by Mn and Fe oxides and diminishes the remaining metal phases. At a higher soil pH, organically complexed Zn predominates and metal associates more with Fe than Mn oxides. In this study, since pH values of all biosolids were adjusted to near neutrality before use, any difference may be attributed to the biosolid matrix, and not the pH of the medium: MOS is physico-chemical biosolid, while BES is biological biosolid. Additionally, although no measurements of metal activity were currently performed, since this was beyond the scope of the present work, we agree with the statement of McBride [38]. He specified that one has to consider the metal activity in soil solution, rather than the total metal concentration, to judge for phytotoxicity of metals. The statement of McBride [38] may serve as a further argument to explain the differences reported above, when comparing our results to those of Li et al. [49], notably with regards to the concentration of metals in water leaching from biosolid amended soils.

## 5 CONCLUSION

The purpose of this glasshouse study was to investigate the effects of spiked Cd, Cu, Zn and metals mixture versus non-spiked MOS and BES biosolids which were non-cleaned, cleaned, and washed on the corn growth as well as on the metals biosorption and leaching. Metal concentrations in plants were quantified in the four separate parts of corn at harvest: roots, stalks, leaves and seeds.

The biosolids investigated in this project contained levels of nutrients suitable for agricultural use despite having undergone cleanup and washing procedures. Micro-nutrients, such as Fe and S, may be supplied to soils from cleaned biosolid, because the leaching process requires Fe- and S-containing chemicals. Results did not show a negative effect of biosolid spiking, cleanup and/or washing on the seed germination and growth parameters of corn. In comparison to SOIL and to non-cleaned biosolid, cleaned biosolid generally enhanced the aerial parts/roots ratios. Furthermore, Cd was not detected in the harvested seeds and stalks, while the Cu level was very low. Migration of Cd and Cu followed a decreasing trend: roots > leaves > stalks, while that of Zn diminished in the order: leaves > roots > stalks. The uptake and biosorption of tested metals were affected by their available fractions, which in turn, were influenced by spiking and cleanup-washing. The remaining elements drifted differently in corn parts, and effects of biosolid cleanup and/or washing were mostly related to biosolid spiked with Cd or Cu. Independently of biosolid amendments, leaching of Cu and Zn into leachate generally diminished with time, while that of Cd was more randomized. Statistical analysis of the concentrations of metals in leachate showed that most of the changes occurred because of cleanup of the MOS biosolid, while few variations were noted with BES biosolid. Concentrations of measured metals in corn tissues and in leachate were generally below their respective critical limits.

Spiking biosolid with a given metal, either alone or in the metal mixture, led to a storage of this element in at least one of the corn parts and/or in leachate, but biosolid cleanup caused a significant decrease in this trend. Interestingly, the restriction imposed by the guidelines of the Quebec government regarding the agricultural spreading of BES biosolid, due to its high Cu content can be avoided if this biosolid is cleaned. In addition, knowing that several soils in the province of Quebec may contain high levels of heavy metals, the use of cleaned instead of non-cleaned biosolid should minimize, or at least delay, the risk of soil enrichment by these metals.

Finally, dissimilarities exist in the ability of MOS and BES biosolids to be cleaned and/or washed by the leaching process. These dissimilarities indicate that the proposed cleanup process depends not only on the initial total metallic charge of biosolid, but also on the biosolid type. Only MOS biosolid washing had few supplemental significant effects on the biosorption and leaching of metals. The present work should be completed by further agri-environmental investigations in glasshouse and field to compare the short-term and long-term fates of biosolid-borne metals following the spreading of biosolid cleaned by our leaching process. Such studies should focus on the measurement of the ion activity of a given metal instead of its total concentration.

## CRediT authorship contribution statement

**Barraoui D**. contributed to the conception and design of the study and the implementation of the experimental design in the greenhouse. He also collected data and wrote the first draft of the manuscript.

**Blais JF**. is the corresponding author and co-PI. He contributed to the conception and design of the study, collaborated to the interpretation of the results. Revised the final version of the manuscript and did the last revision for the submission.

**Labrecque M**. was one of the PI involved in the conception and design of the study. He collaborated to the interpretation of the results and to the revision of the manuscript.

## Acknowledgements

This research has been realized thanks to the valuable aid of the Canada Research Chair Program (grant 950-202886). We are grateful to Gabriel Teodorescu, from the Montreal Botanical Garden for his contribution to the management of this work, to Myriam Chartier and Danielle Leblanc for their technical assistance, and to Stéphane Daigle for his participation to statistical calculations.

